# Synaptic vesicle proteins and ATG9A self-organize in distinct vesicle phases within synapsin condensates

**DOI:** 10.1101/2022.06.14.496103

**Authors:** Daehun Park, Yumei Wu, Aaron Baublis, Pietro De Camilli

## Abstract

Ectopic expression in fibroblasts of synapsin 1 and synaptophysin is sufficient to generate condensates of vesicles highly reminiscent of synaptic vesicle (SV) clusters and with liquid-like properties. We show that unlike synaptophysin, other major integral SV membrane proteins fail to form condensates with synapsin, but coassemble into the clusters formed by synaptophysin and synapsin in this ectopic expression system. Another vesicle membrane protein, ATG9A, undergoes activity-dependent exo-endocytosis at synapses, raising questions about the relation of ATG9A traffic to the traffic of SVs. We have found that both in fibroblasts and in nerve terminals ATG9A does not coassemble into synaptophysin-positive vesicle condensates but localizes on a distinct class of vesicles that also assembles with synapsin but into a distinct phase. Our findings suggest that ATG9A undergoes differential sorting relative to SV proteins and also point to a dual role of synapsin in controlling clustering at synapses of SVs and ATG9A vesicles.

## Main

A defining characteristic of presynaptic sites in the nervous system is the presence of clusters of neurotransmitter containing synaptic vesicles (SVs) anchored to the presynaptic plasma membrane^1^. Upon nerve terminal depolarization, SVs close to the plasma membrane undergo exocytosis to release their neurotransmitter content^2^. Their membranes are then rapidly recaptured by endocytosis and reused for the generation of new SVs that re-enter the vesicle cluster^3-6^. Recent studies have implicated liquid-liquid phase separation as an organizing principle in the assembly of these clusters and have pointed to the protein synapsin as to a key player in these assemblies, also referred henceforth as condensates^7-9^. Synapsin, a dimeric protein associated with SVs by low affinity charge-based interactions, has the intrinsic property to phase separate and captures SVs into its phase^9-11^. Loss of synapsin function by genetic perturbation or by antibody injection results in the dispersion of SVs with the exception of those directly tethered the “active zones” of secretion^9, 12, 13^. Conversely, synapsin can induce vesicle clusters when expressed together with the major SV protein synaptophysin in fibroblasts^14^, cells which physiologically do not express either protein. When expressed alone in this exogenous system, synaptophysin is targeted to small sparse vesicles that can be labeled by endocytic tracers^15, 16^. Upon coexpression with synapsin these vesicles become assembled in large clusters that are morphologically very similar to those found in presynaptic nerve terminals and that share with these clusters liquid properties^14^. Interestingly, several other SV proteins tested do not induce the formation of SV-like clusters when co-expressed with synapsin^14^. These results open the possibility of exploiting this experimental model to gain further insight into SV biogenesis and into the traffic of SV proteins relative to other proteins.

An open question in the cell biology of the presynapse is the relation between the traffic of SV proteins and the traffic of ATG9A. ATG9A is a lipid scramblase^17-19^ required for the growth and expansion of the autophagic membrane and the only transmembrane protein of the core autophagy machinery^20, 21^. ATG9A is present in axon terminals where, together with other autophagy factors, it plays an important role in synapse development and homeostasis^22-24^. Proteomic studies have detected ATG9A (at low copy number) in purified SV fractions^25-27^. Moreover, recent imaging studies have shown that synaptically-localized ATG9A undergoes exo-endocytosis in response to activity in parallel with bona fide SV proteins^22, 24^. However, how its traffic in nerve terminals relates to that of SVs remains elusive.

Here we have used the vesicle cluster reconstitution system in non-neuronal cells to gain insight into the traffic of ATG9A relative to the traffic of SV proteins. More specifically, we investigated which exogeneous proteins can be recruited to the synaptophysin and synapsin positive vesicle condensates. We have found that all of several bona fide SV proteins tested accumulate into the vesicles of these vesicle assemblies. However, we also found that ATG9A does not assemble into them. ATG9A is a component of a distinct set of slightly larger vesicles that can also be clustered by synapsin. Moreover, when both synaptophysin and ATG9A are co-expressed with synapsin, synaptophysin vesicles and ATG9A vesicles segregate in different sub-phases within the synapsin phase. Proteomic analysis further showed a clear distinct composition of the synaptophysin and ATG9A vesicles in this experimental model. A segregation of synaptophysin and ATG9A within nerve terminals was also observed when fluorescently tagged ATG9A and synaptophysin-HA were expressed in neurons. Collectively, our studies suggest a differential sorting of SV proteins and ATG9A in axon endings and the property of synapsin to capture both bona fide SVs and ATG9A vesicles in proximity of presynaptic sites.

## Results

### Synaptophysin and synapsin clusters in fibroblasts recruit other SV components

As we described previously^14^, coexpression of synaptophysin (Syph) and of mCherry-synapsin (mCherry-Syn) in COS7 cells results in the formation of clusters of small vesicles that are similar in size to SVs (Fig. 1a,b). Moreover, like bona fide SVs, the small vesicles can be labeled by the extracellular tracer cholera toxin-horseradish peroxidase (CTX-HRP), revealing and endocytic origin^14^ (Fig. 1b). These vesicles clusters (also referred to as vesicle condensates), appear as large droplets sparse throughout the cytoplasm in fluorescence microscopy (Fig. 1a). Co-expression with synapsin of several other SV proteins^28^ such as secretory carrier membrane 5 (EGFP-SCAMP5), the vesicular glutamate transporter (vGlut fuse to pHluorin, vGlut-pH) and synaptotagmin 1 (SYT1-EGFP) did not results in the formation of similar clusters^14^ (Fig. 1a). SCAMP5 and vGlut localized to sparse vesicles and SYT1 had a predominant plasma membrane localization, as demonstrated in several previous studies^14, 29^ (Fig. 1a). We confirmed these studies and additionally tested other SV proteins^28^: vesicle-associated membrane 2 (VAMP2-pHluorin, VAMP2-pH), the vesicular GABA transporter (vGAT-pHluorin, vGAT-pH) and Rab3A (EGFP-Rab3A). We found that none of them formed condensates with synapsin, which had a diffuse cytosolic distribution irrespective of the co-expression of these proteins (Fig. 1a). The transferrin receptor (transferrin- pHluorin), a protein that undergoes exo-endocytic recycling and that was previously shown to colocalize with synaptophysin when this is expressed in fibroblasts^15^, also did not form condensates with synapsin (Fig. 1a).

**Fig. 1.**
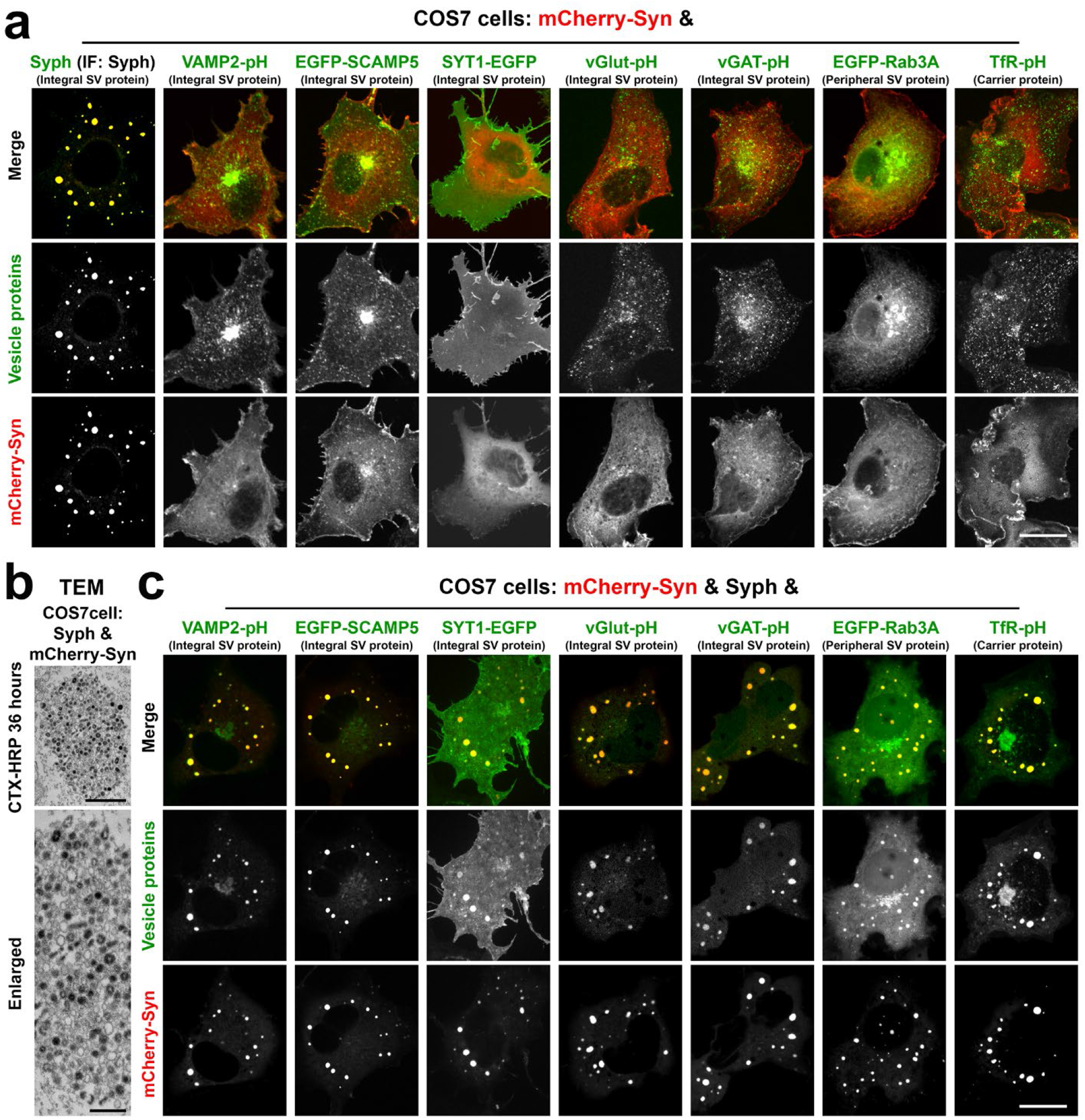
Synaptophysin and synapsin condensates recruit other SV proteins. a, COS7 cells were co-transfected with mCherry-synapsin (Syn) and one of these proteins as indicated: synaptophysin (Syph) (untagged), VAMP2-pHluorin (VAMP2-pH), EGFP-SCAMP5, synaptotagmin 1-EGFP (SYT1-EGFP), vesicular glutamate transporter 1-pHluorin (vGlut-pH), vesicular GABA transporter-pHluorin (vGAT-pH), EGFP-Rab3A and transferrin receptor-pHluorin (TfR- pH). Synaptophysin, which was untagged, was revealed by immunofluorescence. Only cells that co-express mCherry- synapsin and synaptophysin show large droplets. b, Synaptophysin and mCherry-synapsin expressing COS7 cells were exposed to 10 μg/ml cholera toxin conjugated HRP (CTX-HRP) for 36 hours, then fixed, processed for HRP reactivity and embedded for transmission electron microscopy (TEM). Black vesicles are the vesicles labeled by the endocytic tracer CTX-HRP. c, COS7 cells were triple transfected with synaptophysin, mCherry-synapsin and one other fluorescent fusion protein as in field a. Note that all SV proteins and the transferrin receptor coassemble into the droplets formed by synaptophysin and mCherry-synapsin. See also Extended Data Fig. 1. Scale bars, a = 20 μm, b = 500 mm (top) and 200 nm (bottom), c = 20 μm.

However, when these other SV proteins were expressed in COS7 cells together with synaptophysin and with synapsin, all of them co-assembled into the synaptophysin/synapsin condensates (Fig. 1c and Extended Data Fig. 1a-c). Thus, synaptophysin helps nucleate vesicles into which other SV proteins are sorted and which assemble into condensates with synapsin via the low affinity, primarily charge-based, synaptophysin-synapsin interaction^14^. Synaptophysin vesicles may represent the expansion of a physiologically occurring endocytic vesicular compartment, as they are also positive for the transferrin receptor and have an acidic lumen - a feature of house-keeping endocytic vesicles - as shown by alkalinization with NH_4_Cl of cells expressing VAMP2-pH (Extended Data Fig. 1d,e).

### The liquid nature of vesicle clusters in fibroblasts mimics a property of SV clusters in nerve terminals

We previously showed that condensates of synapsin and synaptophysin have liquid-like property^14^. A similar characteristic of the clusters of synaptophysin, other SV membrane proteins and synapsin was demonstrated by their property to fuse with each other and to undergo fluorescence recovery after photobleaching, as exemplified by droplets comprising EGFP-SCAMP5, mCherry-synapsin and synaptophysin (Fig. 2a-d). Not surprisingly, recovery of mCherry-synapsin, a peripheral membrane protein, is much faster than the recovery of an integral vesicle protein, EGFP-SCAMP5, which reflects vesicle exchange between droplets (Fig. 2b-d). Further supporting the role of phase separation mechanisms mediated by synapsin in the formation of the condensates, the droplets containing VAMP2-pH, synaptophysin and mCherry-synapsin were reversibly dissolved by 1,6- Hexanediol (Fig. 2e; Supplementary Video 1).

**Fig. 2.**
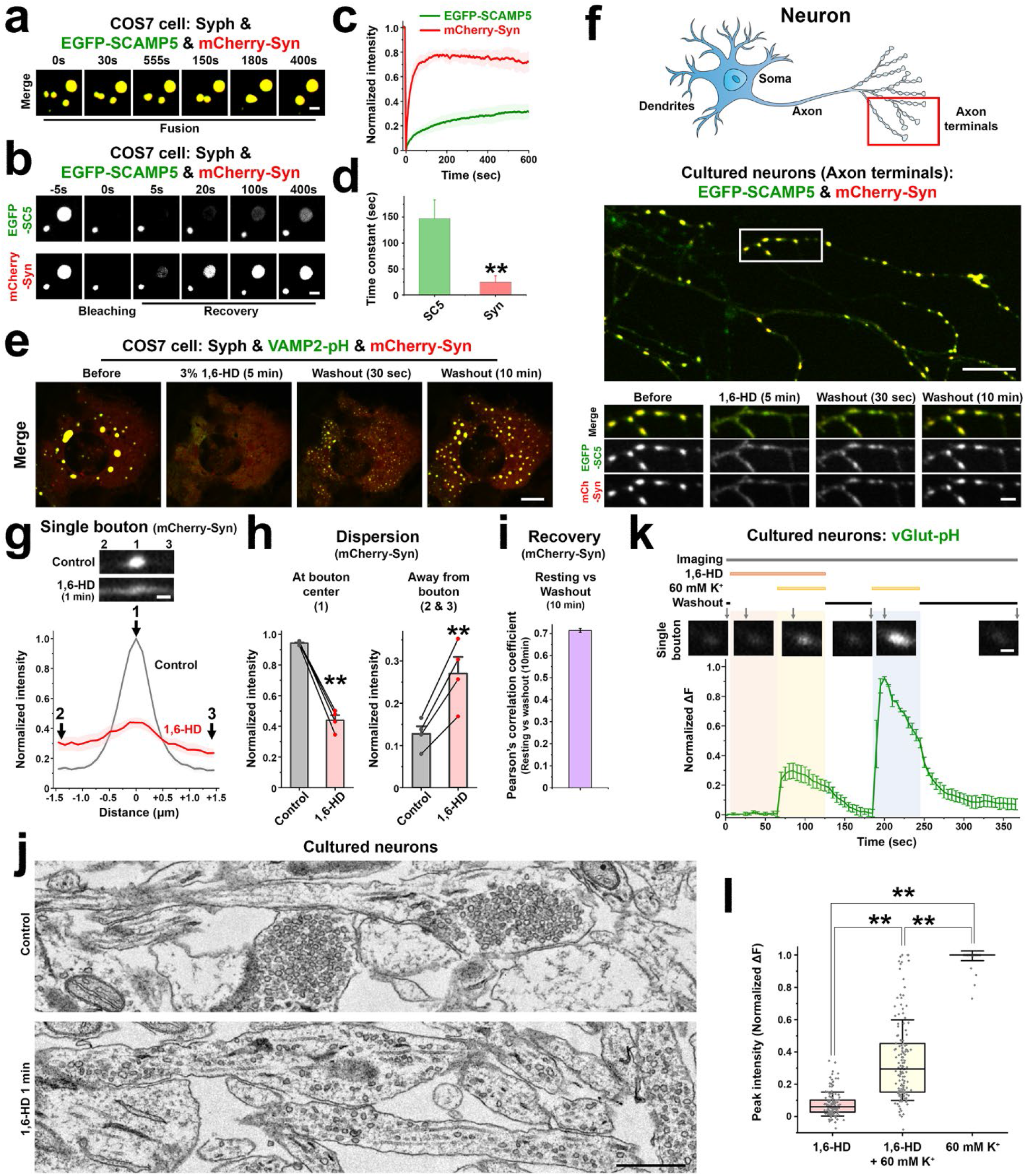
Liquid-like properties of vesicle clusters in fibroblasts and of SV clusters in cultured neurons. a-d, COS7 cells triple transfected with mCherry-synapsin, synaptophysin (untagged) and EGFP-SCAMP5. a, Fusion of droplets. b, Representative time-lapse images showing fluorescence recovery of EGFP-SCAMP5 and mCherry-synapsin after photobleaching of a single droplet. c, Plots of the average fluorescence intensities after photobleaching of multiple EGFP-SCAMP5 and mCherry-synapsin positive droplets. d, Fluorescence recovery traces were fitted to measure the time constant of fluorescence recovery kinetics. Data are represented as mean ± SD (n = 5). e, Droplets comprising VAMP2-pH, mCherry-synapsin and synaptophysin (not shown) in COS7 cells disperse reversibly upon 3% 1,6- Hexanediol (1,6-HD) treatment. See also Supplementary Video 1. f, Live fluorescence imaging of hippocampal neuronal cultures expressing EGFP-SCAMP5 and mCherry-synapsin, treated with 3% 1,6-Hexanediol (1,6-HD) for 5 min and then washed. The cartoon at the top indicates the region of the neuron examined. The micrographs at the bottom show the area bracketed by a rectangle in the main field at high magnification and at different times. Note that SV clusters were reversibly dispersed by 1,6-Hexanediol and that SCAMP5 and synapsin underwent parallel dispersion and re-clustering. See also Supplementary Video 2. g, mCherry synapsin intensity on a presynaptic bouton and flanking axonal regions (± 1.45 μm from each bouton centers) before and 1 min after addition of 1,6-Hexanediol. Fluorescent images are shown at the top. Numbers indicate corresponding regions. h, Statistical comparison of mCherry-synapsin fluorescence intensities at the regions indicated by numbers in g. Values are means ± SEM; \*\**p* < 0.01 by paired t-test (n = 45 boutons from 4 independent experiments). i, The localization of mCherry-synapsin before 1,6-Hexanediol treatment and after 10 min washout was analyzed by calculating the Pearson’s coefficient. Values are means ± SEM (n = 3 independent experiments). j, Control and 1,6-Hexanediol treated (1 min) neuronal cultures were fixed for transmission electron microscopy (TEM). See also Extended Data Fig. 2. k, Normalized average vGlut- pHluorin fluorescence intensity profile and representative time lapse images of presynaptic boutons from cultured hippocampal neurons stimulated with high K^+^ in the presence or absence of 1,6-Hexanediol. See also Supplementary Video 3. Values are means ± SEM. l, Peak vGlut-pHluorin fluorescence intensity values of presynaptic boutons in neurons exposed to 1,6-Hexanediol before and during a first high K^+^ stimulation and then during a second high K+ stimulation in the absence of 1,6-Hexanediol. Box plots show the median line (midline), 25/75 percentiles (boxes), and SD (whiskers) at each condition. \*\**p* < 0.01 by Tukey’s HSD post hoc test (143 boutons from 3 independent experiments were analyzed). Scale bars, a and b = 2 μm, e = 10 μm. f = 10 μm and 2 μm (bottom), g = 1 μm, j = 500 nm, k = 1 μm.

Our hypothesis that vesicle clusters in fibroblasts represent a model of bona fide SV clusters at synapses implies that bona fide SVs in nerve terminals must exhibit the same striking response to 1,6-Hexanediol. This was never tested so far, although the liquid nature of SV clusters was inferred indirectly from the many known properties of such clusters^7^. To address this question, we cultured mouse hippocampal neurons expressing EGFP-SCAMP5 and mCherry-synapsin. Under resting conditions, the fluorescence of the two proteins had the typical (and identical) punctate localization in peripheral axonal branches that reflects presynaptic SV clusters^30^ (Fig. 2f). Similar to what we had observed in fibroblasts (Fig. 2e; Supplementary Video 1), these puncta rapidly (within 1 min) dispersed after addition of 1,6-Hexanediol (Fig. 2f-h; Supplementary Video 2), indicating spreading of SVs along the axon. After washout, they rapidly reassembled at sites of pre-existing clusters (Fig. 2f,i). The conclusion that the 1,6-Hexanediol dependent dispersion of the EGFP- SCAMP5 and mCherry-synapsin fluorescence in nerve terminals reflected the dispersion of SVs was confirmed by electron microscopy (Fig. 2j; Extended Data Fig. 2)

The dispersed and clustered state of SVs correlated with a strong difference in exocytosis in response to depolarization. The secretory response, as assessed by vGlut-pHluorin fluorescence in neurons expressing vGlut-pHluorin and mCherry-synapsin, was much higher when SVs were clustered at presynaptic sites than during exposure to 1,6-Hexanediol, when they were dispersed (Fig. 2k,l; Supplementary Video 3).

### Condensates of ATG9A-positive vesicles upon co-expression of ATG9A with synapsin

We next investigated the relation of ATG9A to SVs. As previously described^31, 32^, when expressed alone, ATG9A-EGFP was enriched in the Golgi complex, but also localized in scattered puncta throughout the cytoplasm (Fig. 3a). Surprisingly, when co-expressed together with mCherry-synapsin, ATG9A formed large droplets (Fig. 3b) similar to those found in synaptophysin and mCherry-synapsin expressing cells (Fig. 1a). Correlative light-electron microscopy (CLEM) further revealed that ATG9A-EGFP and mCherry-synapsin droplets represented clusters of small vesicles (Fig. 3c) that resemble the synaptophysin and mCherry-synapsin condensates (Fig. 1b), although the ATG9A vesicles were slightly larger [diameter: 41.78 nm (synaptophysin) vs 54.88 nm (ATG9A)] (Fig. 3d). As in the case of synaptophysin vesicles, the vesicles in ATG9A-EGFP and mCherry-synapsin condensates could be labeled by the endocytic tracer CTX-HRP (Fig. 3e), indicating that they are part of the endocytic system. Moreover, CTX-HRP labeled ATG9A-EGFP vesicles intermixed at random with unlabeled vesicles consistent with a liquid like nature of the vesicle clusters (Fig. 3e). Thus, similarly to synaptophysin, but in contrast to other SV proteins tested, ATG9A, when expressed in fibroblasts without synaptophysin accumulated in vesicles that could be trapped into liquid condensates by an interaction with synapsin. We had shown that a feature of synaptophysin that was at least in part responsible for its property to co-assemble with synapsin was its negatively charged cytoplasmic tail, which establishes charge-based interactions with the highly positively charged C-terminal region of synapsin^14^. Interestingly, ATG9A also has a negatively charged cytosolic tail which could establish electrostatic interactions with synapsin (Fig. 3f).

**Fig. 3.**
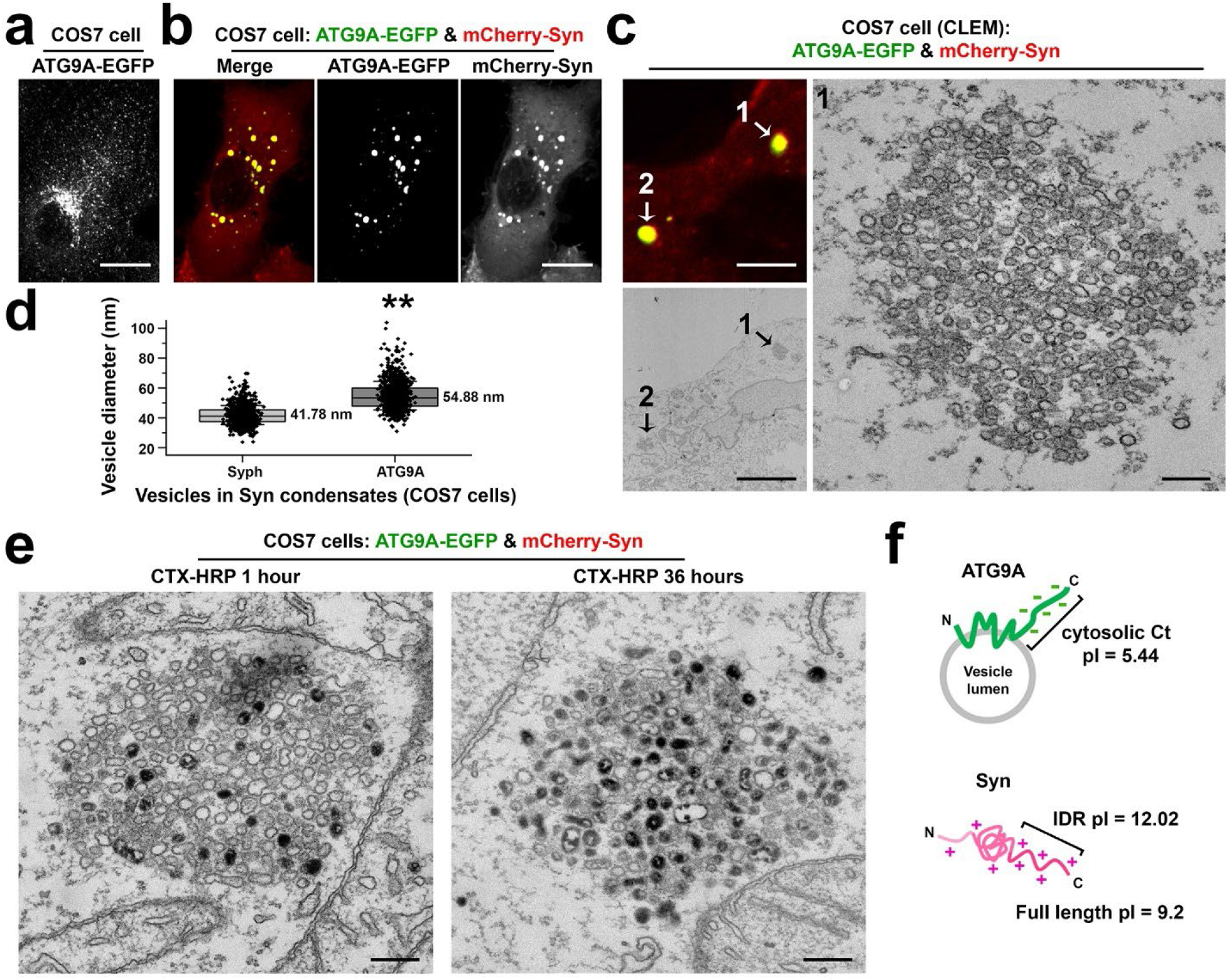
Co-expression of ATG9A-EGFP with mCherry-synapsin results in clusters of small vesicles. a,b, ATG9A- EGFP was expressed either alone (a) or with mCherry-synapsin (b) in COS7 cells. c, Correlative light-electron microscopy (CLEM) of a COS7 cell co-expressing ATG9A-EGFP and mCherry-synapsin. Left: arrowheads point to droplets positive for ATG9A-EGFP and mCherry-synapsin in the fluorescence and EM images. Right: high magnification EM image of droplet #1. d, Size distribution of the ATG9A and synaptophysin vesicles when these two proteins are expressed independently with synapsin. Box plots show the median line (midline), 25/75 percentiles (boxes), and SD (whiskers). Data are represented as mean ± SD. 1,000 vesicles were measured for each group (from three independent experiments). \*\**p* < 0.01, by Student’s t test. e, COS7 cells co-expressing ATG9A-EGFP and mCherry-synapsin were treated with 10 μg/ml cholera toxin conjugated HRP (CTX-HRP) for either 1 hour (left) or 36 hours (right) at 37°C and then fixed for transmission electron microscopy (TEM). f, Schematic drawing of synapsin and ATG9A. pI = isoelectric point. IDR: intrinsically disordered region (C-terminal region of synapsin). Scale bars, a and b = 20 μm, c = 5 μm (left) or 200 nm (right), e = 200 nm.

### ATG9A and synaptophysin form two distinct vesicle clusters within synapsin condensates

Next, we expressed in fibroblasts ATG9A-EGFP together with synaptophysin and with mCherry-synapsin. Strikingly, ATG9A and synaptophysin segregated in distinct phases within the mCherry-synapsin phase, with synapsin being more concentrated in the synaptophysin subphase (Fig. 4a-d). CLEM analysis of cells previously exposed to CTX-HRP for 36 hours further showed that both phases were represented by clusters of small vesicles labeled by the extracellular tracer (Fig. 4e). However, the vesicles in the ATG9A-EGFP/mCherry-synapsin phase were slightly bigger than the vesicles in synaptophysin/mCherry-synapsin phase, consistent with the different sizes of the two sets of vesicles when ATG9A and synaptophysin are expressed independently (Fig. 3d, 4e and Extended Data Fig. 3). Both components of ATG9A/synaptophysin/synapsin condensates had liquid-like properties, as shown by their property to undergo reversible dispersion by 1,6- Hexanediol (Fig. 4f; Supplementary Video 4), to fuse (Fig. 4g) and to undergo fluorescence recovery after photobleaching (Fig. 4h).

**Fig. 4.**
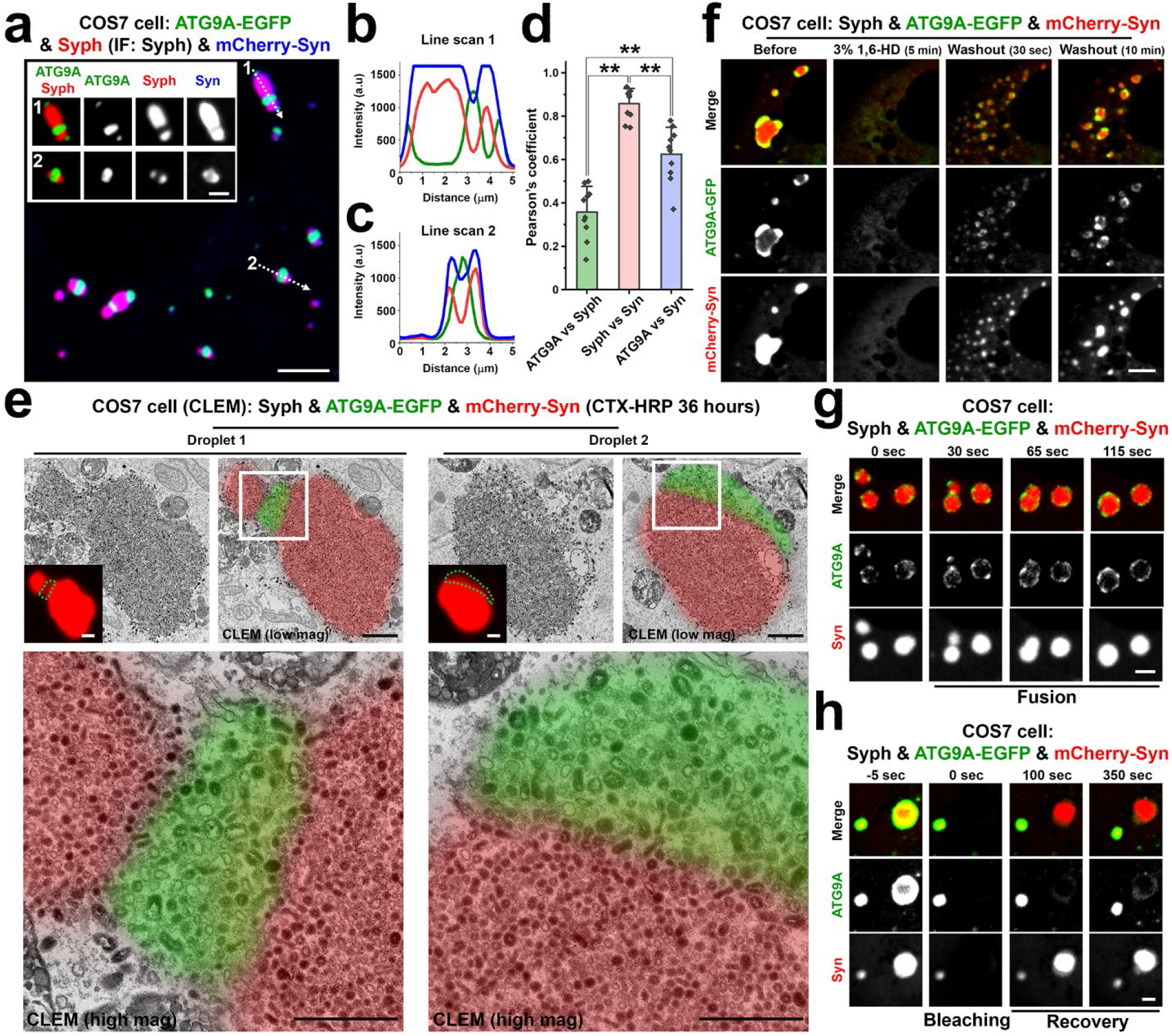
Synaptophysin and ATG9A segregate in distinct vesicle clusters within the synapsin phase. COS7 cells were triple co-transfected with synaptophysin (untagged), ATG9A-EGFP and mCherry-synapsin. a-c, Representative confocal image (a), and corresponding line-scan analysis of the droplets (b,c) showing that synapsin phases include two distinct sub-phases containing synaptophysin and ATG9A-EGFP, respectively. Synaptophysin was visualized by immunofluorescence. The intensity of the mCherry-Synapsin fluorescence is more intense on the synaptophysin phase, as quantified in d. d, The colocalization of the pairs of proteins indicated was calculated by Pearson’s coefficients. Values are means ± SD. \*\**p* < 0.01 by Tukey’s HSD post hoc test (n = 10 independent experiments). e, Synaptophysin, ATG9A-EGFP and mCherry-synapsin co-expressing COS7 cells were incubated with 10 μg/ml cholera toxin conjugated HRP (CTX-HRP) for 36 hours and then fixed for correlative light-electron microscopy (CLEM). Green dotted lines indicate ATG9A-EGFP positive regions in mCherry-synapsin condensates. Note that synaptophysin and ATG9A vesicles have slightly different sizes. See also Extended Data Fig. 3. f-h, Dynamics of the ATG9A, synaptophysin and synapsin droplets: f, Reversible dispersion by 1,6-Hexanediol treatment. See also Supplementary Video 4; g, Fusion; h, Fluorescence recovery after photobleaching. Scale bars, a = 5 μm (2 μm for insets), e = 1 μm (low mag and inset) or 500 nm (high mag), f = 5 μm, g and h = 2 μm.

### Distinct features of synaptophysin vesicles and ATG9A vesicles as revealed by proteomic analysis

The segregation in two distinct sets of vesicles of synaptophysin and exogenous ATG9A in fibroblasts prompted us to perform a proteomic analysis to further characterize their difference. We first used proximity biotinylation, with the rationale that fusing a biotinylation enzyme to a vesicle protein would results in biotinylation of other components of the vesicle given the fluid nature of membranes. To this aim, a construct encoding miniTurboID (mTB), an enzyme that allows rapid (within minutes) labeling of neighboring proteins within a ∼10 nm range^33^, was fused in frame to the cytosolic C-terminus of synaptophysin and of ATG9A, followed by an HA tag and expressed in COS7 cells. An additional construct encoding an N-terminal ER targeting sequence followed by miniTurboID (expected to direct the localization of miniTurboID at the ER surface^33^) was used as a control in initial experiments. Anti-HA tag immunofluorescence revealed that fusion of ATG9A and synaptophysin to miniTurboID-HA did not change the subcellular localization of these two membrane proteins (Extended Data Fig. 4a-c). After15 min incubation with exogenous biotin, biotinylated proteins were detected with streptavidin by either microscopy or western blotting (Extended Data Fig. 4a-c and Fig. 5a,b). Most importantly, the biotinylation signal precisely overlapped with HA-tag immunoreactivity, confirming confinement of the reaction to the neighborhood of the fusion proteins (Extended Data Fig. 4a-c). Biotinylated proteins in extracts of these cells were affinity-purified by streptavidin and three biological replicates for each of the two conditions were analyzed by LC/MS-MS. As illustrated in Fig 5c, which shows the relative enrichment of biotinylated proteins in the extracts of cells expressing ATG9A-miniTurboID-HA and synaptophysin-miniTurboID-HA, respectively, numerous shared proteins were found (gray dots in Fig. 5c). However, substantial differences between the two biotinylated proteomes were also detected. Consistent with previous studies, ATG9A samples were enriched in subunits of AP4 (an heterotetrameric complex responsible for the sorting of ATG9A at the TGN)^31, 32, 34^, in AP4 interactors (tepsin and rusc2)^32, 34, 35^, in subunits of the PIKfyve complex (a phosphoinositide metabolizing complex that that function at late endosomes/lysosomes)^36^ as well as in the autophagy factors ULK1 and ATG13^37^ (Fig. 5c). Conversely, synaptophysin samples were enriched in proteins, or paralogues of proteins, that are found at high concentration in SVs in nerve terminals^26^, such as Rab3, SCAMPs, vacuolar ATPase subunits, VAMP3, Rab27b, as well as other proteins implicated in SV recycling such as Bin1 (amphiphysin2)^38^ and synaptojanin2^39^ (Fig. 5c). Detection of many vacuolar ATPase subunits is consistent with the acidic pH of the lumen of synaptophysin vesicles as mentioned above (Extended Data Fig. 1d,e).

**Fig. 5.**
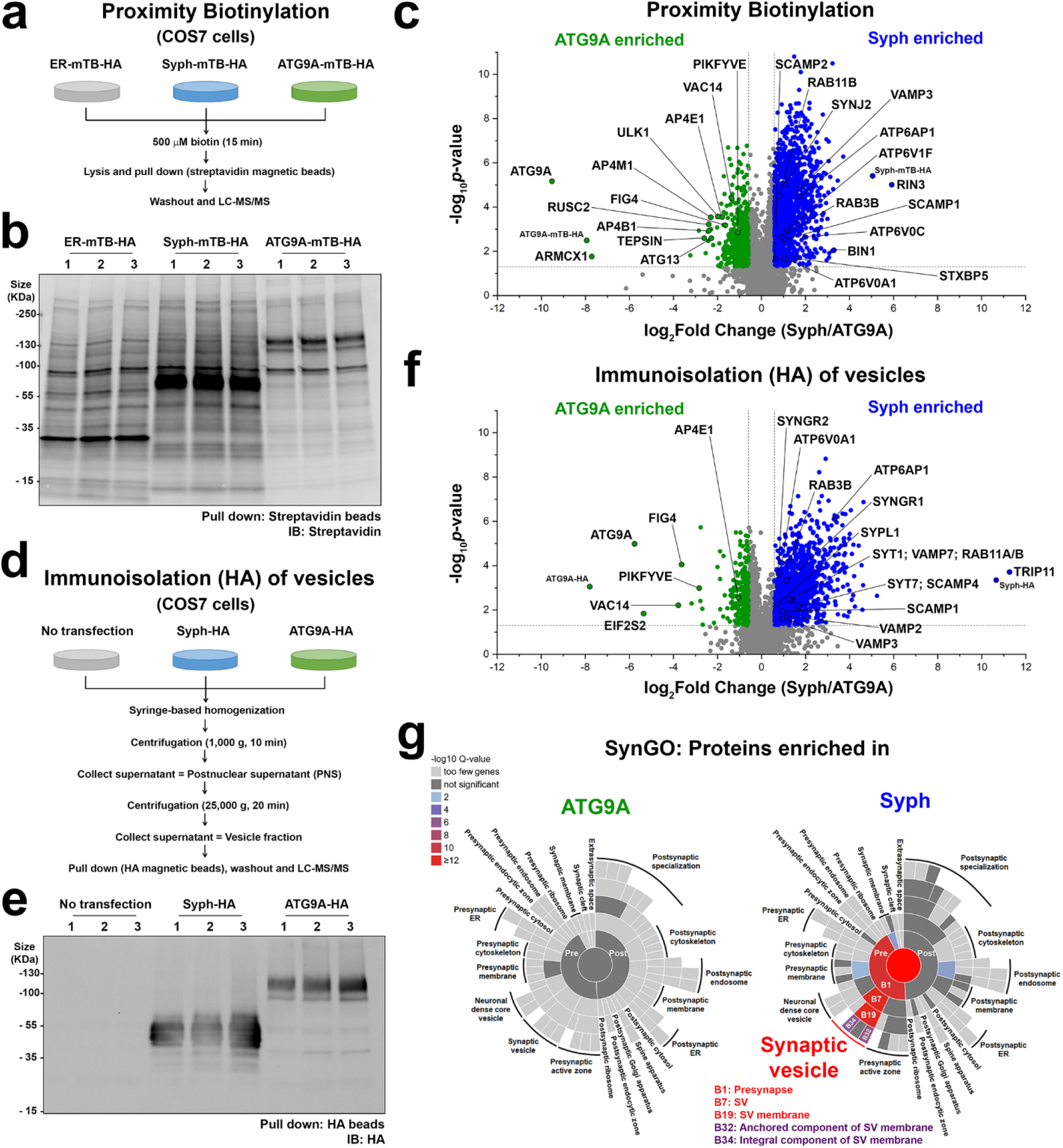
Proteomic analysis of ATG9A and synaptophysin vesicles. a-c, Proximity labeling of synaptophysin and ATG9A vesicles using miniTurboID (mTB). a, Experimental design. b, Biotinylated proteins were affinity-purified on streptavidin magnetic beads and detected by SDS-PAGE and blotting with IRDye680-streptavidin. See also Extended Data Fig. 4a-c. c, Volcano plot of the proteins identified by LC-MS/MS. Significant hits (p-value < 0.05 and absolute fold change > 1.5) are indicated in green (ATG9A enriched) or blue (synaptophysin enriched). d-f, Immunoisolation of synaptophysin-HA and ATG9A-HA vesicles on anti-HA magnetic beads. d, Experimental design. e, A small vesicle enriched cell fraction was immunoisolated on anti-HA magnetic beads. Then proteins were solubilized and processed by SDS-PAGE and immunoblotting with anti-HA antibodies. See also Extended Data Fig. 4d-f. f, A volcano plot showing the proteins enriched in ATG9A-HA or synaptophysin-HA vesicles. Significant hits (p-value < 0.05 and absolute fold change > 1.5) are indicated in green (ATG9A enriched) or blue (synaptophysin enriched). g, Ontology Analysis of the proteome using the SynGO tool showing enrichment of SV proteins in synaptophysin-HA but not in ATG9A-HA immunoisolated samples. Colors represent enrichment Q value (-log10 values). Quantitative LC-MS/MS data from biological triplicates. See also Extended Data Fig. 5.

To complement results of proximity biotinylation, we next performed immunoisolation of the two sets of vesicles from extracts of COS7 cells expressing either synaptophysin-HA or ATG9A-HA. 25,000 g supernatants were used as starting material for these immunoisolations to minimize contamination of large organelles (Fig 5d-f and Extended Data Fig. 4d-f). Quantitative LC-MS/MS of the affinity-purified vesicles (Fig 5f) yielded results that aligned with those of proximity labeling (Fig 5c). In spite of the use of fibroblasts as starting material, SynGO^40^ analysis revealed enrichment in the synaptophysin vesicles of SV proteins or their paralogs (synaptogyrins, synaptotagmins, VAMP2, 3 and 7, SYPL1 and vacuolar ATPase subunits) as well as of proteins with a function in the regulation of the SV cycle (Fig. 5g and Extended Data Fig. 5). In contrast, only very few proteins were enriched in the ATG9A vesicles, relative to synaptophysin vesicles. One of them is a PIKfyve complex (Vac14, PIKfyve and Fig4) (Fig. 5f), in agreement with results of proximity biotinylation (Fig. 5c). Results of this proteomics analysis were validated by protein expression studies. Thus, when mCherry-tagged SV proteins (VAMP2, SCAMP5 or Rab3A) were co-expressed with ATG9A-EGFP, synaptophysin and synapsin, they were enriched in the synaptophysin subphase within the synapsin phase (Fig. 6a,b). Conversely, VAC14 was enriched in the ATG9A subphase (Fig. 6b).

**Fig. 6.**
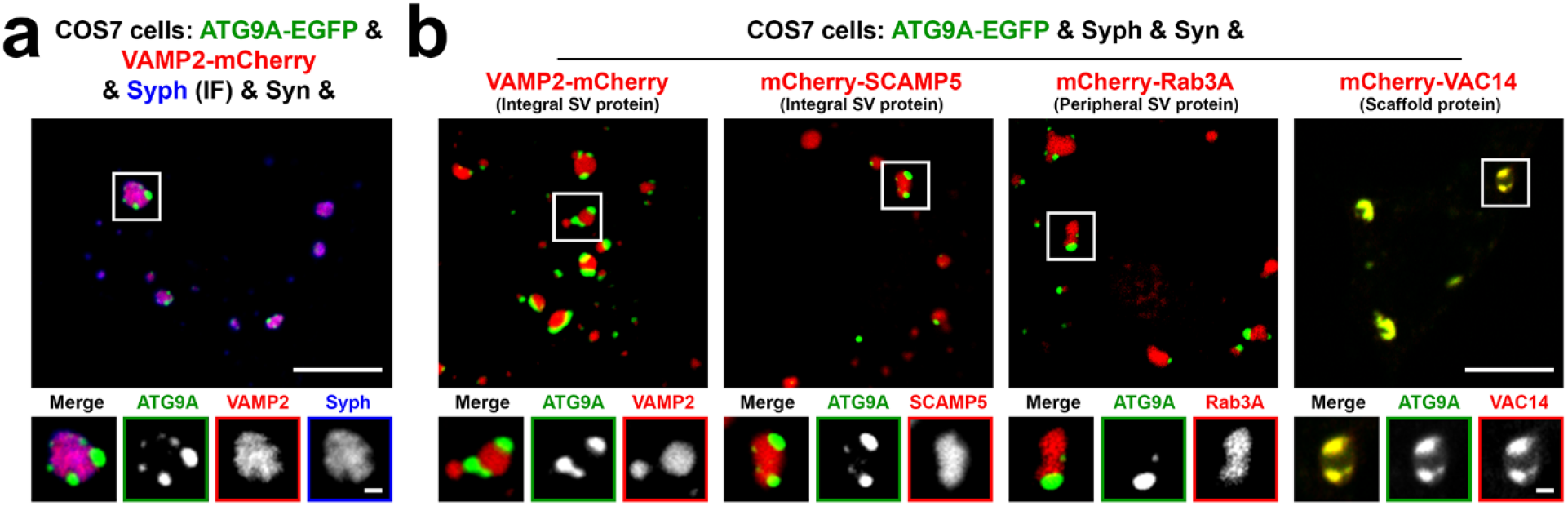
Validation of the proteomic analysis of ATG9A and synaptophysin vesicles. a, Segregation of VAMP2- mCherry from ATG9A-EGFP in COS7 cells also expressing synapsin and synaptophysin. Synaptophysin was visualized by immunofluorescence. b, ATG9A is segregated from SV proteins (VAMP2, SCAMP5, Rab3A) but precisely colocalizes with VAC14 in COS7 cells also expressing synaptophysin and synapsin, consistent with proteomics data. Scale bars, a and b = 10 μm (1 μm for high magnification images).

ATG9A has a conserved YXXØE motif (YQRLE) in its N-terminal cytosolic region (Fig. 7a), which is required for ATG9A exit from the trans-Golgi network (TGN) by mediating capture of ATG9A into AP4 coated buds^31^. Accordingly, AP4 binding defective ATG9A mutant (ATG9A^Y8A^) massively accumulated at the TGN, at a position well separated by the cis-Golgi marker GM130, where it precisely colocalized with the TGN marker TGN46 and failed to form sparse cytoplasmic droplets in the presence of mCherry-synapsin (Fig. 7b; Supplementary Video 5). In fact, when synapsin was co-expressed with ATG9A^Y8A^, it accumulated together with ATG9A^Y8A^ in the TGN area (Fig. 7c), confirming its property to co-assemble with ATG9A. CLEM analysis further revealed that this accumulation of mutant ATG9A and TGN46 correlated with a striking accumulation of small heterogeneously sized vesicles (Fig. 7c).

**Fig. 7.**
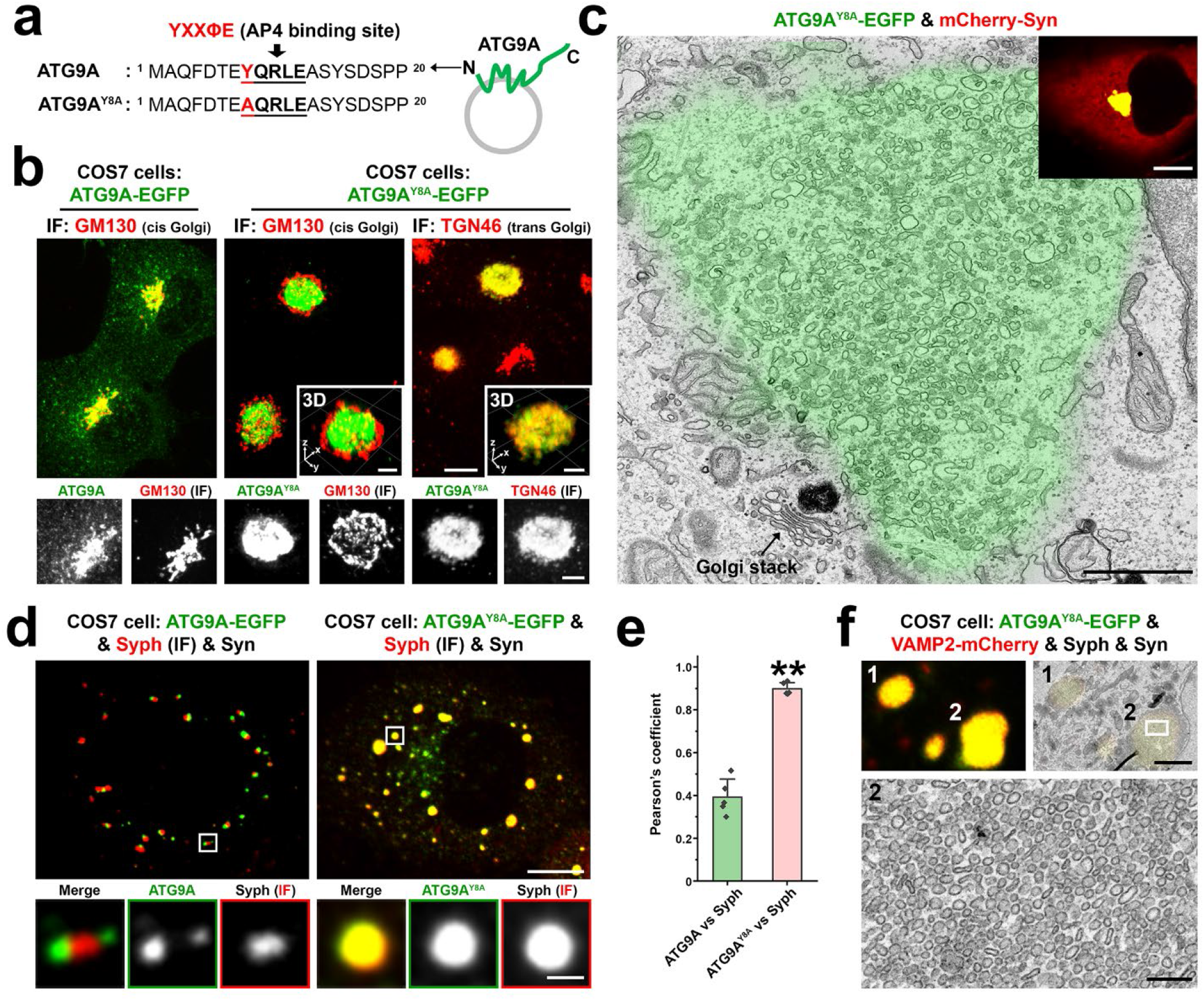
Impact of the Y8A mutation of ATG9A on its sorting and segregation from synaptophysin. a, Amino acid sequence of the cytosolic N-terminal region of ATG9A with the AP4 binding motif in bold characters. b, COS7 cells expressing ATG9A-EGFP (WT) or ATG9A^Y8A^-EGFP and immunostained for cis-Golgi (GM130) and trans-Golgi (TGN46) markers, showing that ATG9A^Y8A^ is retained in the TGN area, where it colocalizes with TGN46. See also Supplementary Video 5. c, Correlative light electron microscopy (CLEM) of a COS7 cell expressing ATG9A^Y8A^-EGFP and mCherry-synapsin, showing that the accumulation of ATG9A^Y8A^ in the TGN area (green color from the fluorescence image) correlates with a massive accumulation of small heterogeneously sized vesicular-tubular structures in the same area. An arrow points to a Golgi stack. The inset shows that synapsin is colocalized with ATG9A^Y8A^ in the TGN area, while no sparse droplets positive for ATG9A^Y8A^ and for synapsin are present in this cell. d, COS7 cell triple transfected with synapsin, synaptophysin and either ATG9A-EGFP or ATG9A^Y8A^-EGFP. Synaptophysin was detected by immunofluorescence. ATG9A^Y8A^-EGFP failed to form distinct clusters within synapsin condensates and completely intermixed with synaptophysin. e, Colocalization analysis of synaptophysin versus either ATG9A (WT) or ATG9A^Y8A^ in COS7 cells also expressing synapsin. Values are means ± SD. \*\**p* < 0.01 by Student’s t test (n = 5 independent experiments). f, CLEM image of a COS7 cell quadruple transfected with ATG9A^Y8A^-EGFP, VAMP2-mCherry, synaptophysin and synapsin. Top images show corresponding light microscopy and EM fields and the bottom image shows a detail of droplet #2 at high magnification. Scale bars, b = 10 μm (5 μm for insets and enlarged images), c = 1 μm (10 μm for inset), d = 10 μm (1 μm for enlarged images), f = 2 μm (200 nm for the enlarged image at the bottom).

Surprisingly, when ATG9A^Y8A^-EGFP was expressed with both mCherry-synapsin and synaptophysin, it was not retained in the TGN and precisely colocalized with synaptophysin and synapsin in droplets (vesicle condensates) sparse around the cytosol (Fig. 7d-f). We suggest that under these conditions ATG9A^Y8A^-EGFP follows a default pathway and that the interaction of ATG9A with AP4 is needed to sort ATG9A away from this pathway.

### Localization of ATG9A in nerve terminals

We next examined the localization of ATG9A-EGFP in nerve terminals when expressed in cultured hippocampal neurons. When ATG9A-EGFP was co-expressed with synaptophysin-HA, the two proteins accumulated at distinct, although juxtaposed, spots in nerve terminals (Fig. 8a,b), On the contrary, when mCherry-SCAMP5 was co-expressed with synaptophysin-HA, these two proteins precisely colocalized at presynaptic sites (Fig. 8a,b). When ATG9A-EGFP, mCherry-synapsin and synaptophysin-HA were co-expressed together, the juxtaposed spots of ATG9A-EGFP and of synaptophysin-HA were both positive for synapsin as we had observed in fibroblasts (Fig. 4a), although, again as in fibroblasts, synapsin was more concentrated in the synaptophysin-region (Fig. 8c).

**Fig. 8.**
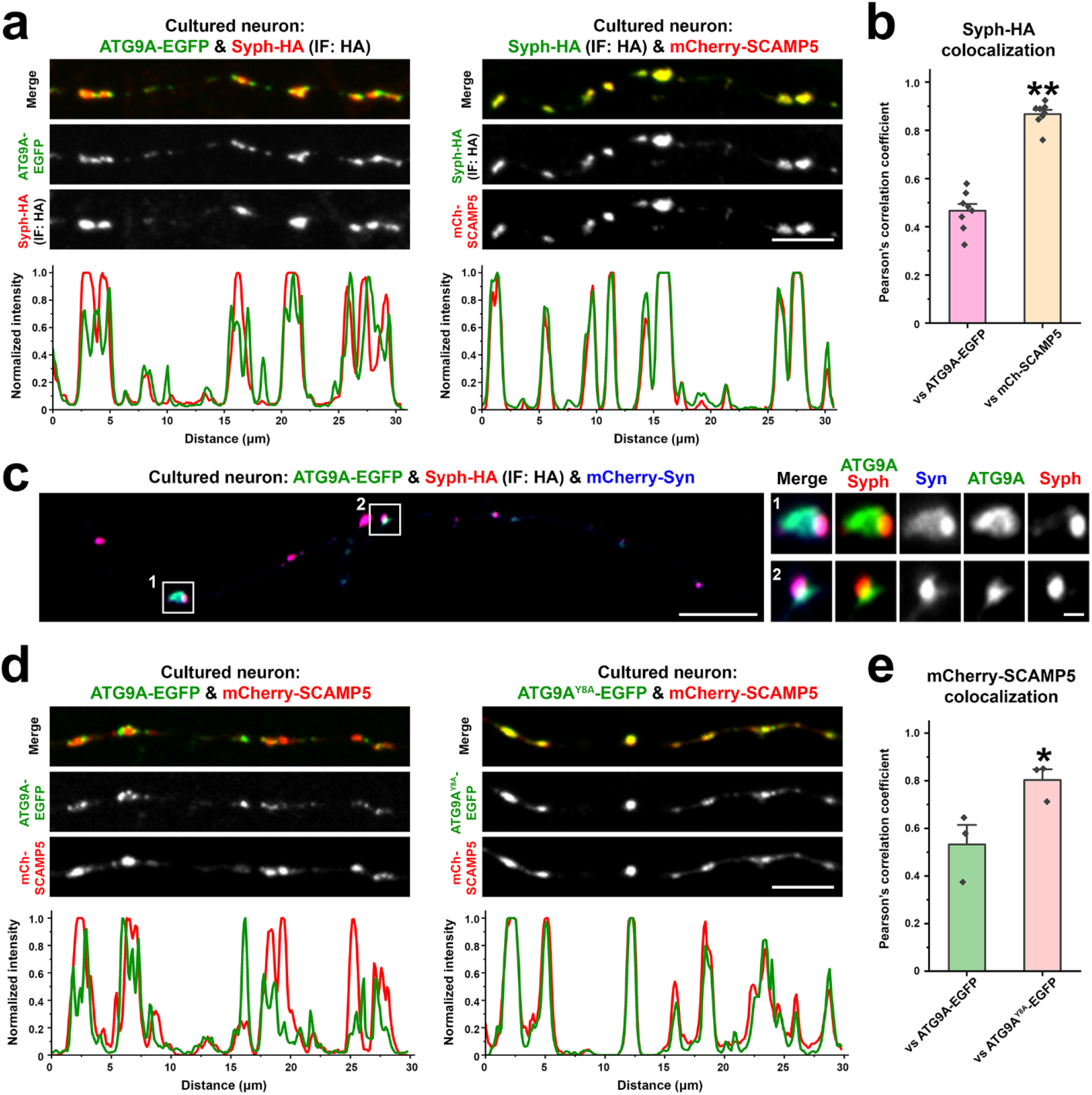
ATG9A vesicle localization in neurons. a-e, Mouse hippocampal neuronal cultures were transfected as indicated and imaged at DIV14-17. a, Representative confocal images (top), and corresponding line-scan analysis (bottom) of neuronal cultures expressing synaptophysin-HA with ATG9A-EGFP or mCherry-SCAMP5. Synaptophysin-HA was detected by anti-HA immunofluorescence. b, Colocalization analysis calculated by Pearson’s coefficients. Values are means ± SEM. \*\**p* < 0.01 by Student’s t test (n = 8 independent experiments). c, Localization of ATG9A-EGFP, synaptophysin-HA and mCherry-synapsin in presynaptic varicosities of an axon. Synaptophysin- HA was visualized by anti-HA immunofluorescence. d, Axonal varicosities of an axon expressing mCherry-SCAMP5 with either ATG9A-EGFP or ATG9A^Y8A^-EGFP. Corresponding line-scans are shown at the bottom. e, Colocalization analysis. Values are means ± SEM. \**p* < 0.05 by Student’s t test (n = 3 independent experiments). Scale bars, a = 5 μm, c = 10 μm (1 μm for high magnification images), d = 5 μm.

The AP4 binding defective mutant of ATG9A (ATG9A^Y8A^) was present in nerve terminals (Fig. 8d) (indicating that in neurons it can exit the TGN, at least under conditions of overexpression) and colocalized with the SV marker, SCAMP5, supporting the importance of AP4 binding for its proper sorting and trafficking in both fibroblasts (Fig. 7d-f) and neurons (Fig. 8d,e).

## Discussion

Results of this study show that the generation of SV-like organelles in fibroblasts is a powerful experimental system to gain insight into mechanisms of SV biogenesis and clustering. We had previously shown that even in this exogenous system SV proteins can self-organize into small vesicles that share several characteristic properties with bona fide SVs such as similarity in size, property to undergo exo-endocytosis and clustering by synapsin into a liquid phase^14^. We now demonstrate that all of several exogenous SV proteins tested coassemble with exogenous synaptophysin in these vesicles and that synaptophysin is the critical driver of these assemblies, at least in this exogenous system. This finding opens the possibility of using this system to gain new insight into the still open question of how SV proteins coassemble into such a specialized vesicle.

We also show that ATG9A accumulates in a different class of slightly larger vesicles, although even such vesicles can be captured in the synapsin phase. The different protein composition of the two types of vesicles, as revealed by a proteomic analysis, further demonstrates that they are part of two distinct vesicle traffic pathways that also diverge in nerve terminals. The list of endogenous proteins that co-enrich with synaptophysin vesicles in COS7 cells supports the hypothesis that SVs represent a special adaptation of a house-keeping vesicle recycling endocytic pathway, as several such proteins are also components of SVs or their paralogues. On the other hand, the enrichment of endogenous proteins implicated in lysosome function (PIKfyve complex^36^) and autophagy (ULK1 complex^37^) in ATG9A vesicles is in line with a role of this protein in protein degradation.

Another important result of our study is that the rapid and reversible dispersion of synapsin-dependent SV-like condensates observed in fibroblasts upon addition of 1,6-Hexanediol is phenocopied in axon terminals. This finding adds a key missing piece of evidence to the concept that native SV clusters are organized according to liquid-liquid phase separation principles. The dispersion-reclustering correlates with a strong difference in depolarization evoked SV induced by high K^+^, consistent with the expectation that SV clustering at presynaptic sites plays a critical role in allowing a sustained secretory response to massive stimulation. Dispersion of active zone proteins^41^ may contribute to the effect of 1,6-Hexanediol on secretion, as phase separation principles were proposed to contribute also to the assembly of many such proteins, including RIM, RIM-BP, ELKS and liprin-α^42-45^.

Presynaptically localized ATG9A was recently shown to undergo activity-dependent exo-endocytosis in parallel with SV proteins in C. elegans^22, 24^. However, consistent with our present findings, even in worms ATG9A did not appear to precisely colocalize with SVs: 1) its localization clearly diverged from the localization of SVs after perturbation of endocytosis^22, 24^ and 2) the localization of SV proteins and of ATG9A was differentially affected by genetic perturbation of the active/periactive zone protein Clarinet^24^. Moreover, while mass spectrometry analysis of purified SV fraction from rat brain detected ATG9A in such vesicles^25-27^, the amount of ATG9A detected was extremely low^26^. It was estimated that 4 copies of ATG9A were present in 100 vesicles^26^. Considering that ATG9A occurs as a trimer, there would be only 1-2 ATG9A trimers per 100 vesicles in presynaptic nerve terminals (versus 525 Synaptophysin hexamers per 100 SVs)^26, 28^. Thus, these findings are consistent with a segregation of ATG9A from bona fide SVs. The concentration of ATG9A in small vesicles is in line with evidence that ATG9A has curvature promoting properties^21^. A propensity to generate or assembly into vesicles larger than bona fide SVs could in principle contribute to its segregation from synaptophysin vesicles. However, we show here that the segregation of ATG9A from synaptophysin occurs only if ATG9A can bind AP4, an adaptor protein shown to be required for the sorting of ATG9A at the TGN. An ATG9A construct lacking the AP4 binding site can surprisingly exit the TGN if co-expressed with synaptophysin, and precisely colocalizes with synaptophysin vesicle clusters both in fibroblasts and in nerve terminals.

The property of ATG9A vesicles to be clustered with synapsin both in fibroblasts and in nerve terminals was unexpected. As discussed above in Results, this property likely relies on the negative charge of the cytosolic C-terminal region of ATG9A, which similarly to the negatively charge C-terminal tail of synaptophysin, may interact with the basic C-terminal region of synapsin. However, ATG9A and synaptophysin vesicles segregate in different subphases within the synapsin phase. Mechanisms that may contribute to this difference may include a different affinity for synapsin, a different concentration of binding sites for synapsin on the ATG9A vesicles or the different diameter of synaptophysin and ATG9A vesicles. It remains possible that the formation of two well distinct vesicle subphases within the synapsin phase may be the consequence of the overexpression of ATG9A, as while ATG9A plays an important role at synapses, there is no evidence for its presence at high concentration in a purified small vesicle fraction from rat brain synaptosomes^26^. Nevertheless, our study demonstrates that, both in nerve terminals and in an exogenous system, major SV proteins and of ATG9A are differentially sorted and represents an important step towards the elucidation of how SV traffic and vesicle traffic responsible for autophagosome formation are interconnected.

## Methods

### Plasmid DNA construction

Human ATG9A was amplified from pMXs-puro-RFP-ATG9A from Addgene (plasmid #60609) and subcloned into the EGFP-N1 vector. ATG9A Y8A-EGFP was generated using site-directed mutagenesis (QuikChange II XL, Agilent Technologies) of ATG9A- EGFP. The following plasmids were previously described: Synaptophysin^14^, mCherry-synapsin^14^, EGFP-SCAMP5^46^, Synaptotagmin1-EGFP^29^, mCherry-Rab3A^47^ and mRFP-Rab5A^47^. Synaptophysin-HA or ATG9A-HA constructs were generated by inserting the HA epitope sequence at the C-terminus of the synaptophysin or ATG9A sequences. miniTurboID was amplified from Addgene (plasmid #107174) and inserted between synaptophysin and the HA tag or between ATG9A and the HA tag to obtain synaptophysin-miniTurboID-HA and ATG9A- miniTurboID-HA. The HA tag and a stop codon were inserted between miniTurboID and the V5 tag of Addgene plasmid #107174 to generate ER-miniTurboID-HA. Rab3A was amplified by PCR from the mCherry-Rab3A^47^ and cloned into the EGFP-C1 vector to construct EGFP-Rab3A. Human VAC14 was obtained from Addgene (Plasmid #47418) and subcloned into the mCherry- C1 vector. The following plasmid were kind gift: Transferrin receptor-pHluorin (from Dr. David Zenisek, Yale University), VAMP2-pHluorin (from Dr. James Rothman, Yale University), vGlut-pHluorin (from Dr. John Rubenstein, University of California San Francisco), vGAT-pHluorin (from Dr. Susan Voglmaier, University of California San Francisco). Each construct was validated by DNA sequencing.

### Antibodies

The following antibodies were used for immunofluorescence: anti-synaptophysin (101 002, Synaptic Systems), anti-HA (G246, Covance), anti-GM130 (610822, BD Bioscience) and anti-TGN46 (AHP500GT, AbD Serotec).

### Cell culture and transfection

COS7 cells were grown in DMEM supplemented with 10% FBS, 100 U/ml penicillin and 100 mg/ml streptomycin. Cells were maintained at 37°C in a 5% CO_2_ humidified incubator and were transfected by Lipofectamine-2000. Hippocampal neurons were cultured from post-natal day 0 newborn mouse pups and transfected at 8 days in vitro (DIV) using a calcium phosphate transfection method. Neurons were used at days in vitro (DIV) 14 to 18. All animal experiments were approved by the Institutional Animal Care and Use Committee of Yale University.

### Correlative light and electron microscopy (CLEM)

COS7 cells were plated on 35 mm gridded, glass-bottom MatTek dish (P35G-1.5-14-CGRD) and transfected as indicated. Cells were fixed with 4% PFA in 0.1 M phosphate buffer (pH7.3) (PB) and washed in PB. Regions of interest were selected by confocal microscopy and their coordinates were identified using phase contrast. Secondary fixation and TEM sample preparation were performed as described previously^14^.

### Fluorescence imaging

Cells were imaged with a spinning disk confocal microscope using a planar Apo objective 60x, 1.49- NA and an EM-CCD camera (C9100-50; Hamamatsu Photonics) under the control of Improvision UltraVIEW VoX system (PerkinElmer). Live cell imaging buffer (Invitrogen) was used for live cell imaging of COS7 cells. For 1,6-Hexanediol experiments, cells were briefly washed in live cell imaging buffer, incubated with 3% 1,6-Hexanediol in live cell imaging buffer and then returned to live cell imaging buffer alone. For live imaging of neurons, neuronal cultures were switched form neurobasal medium to Tyrode buffer (136 mM NaCl, 2.5 mM KCl, 2 mM CaCl_2_, 1.3 mM MgCl_2_, 10 mM HEPES and 10 mM glucose). 3% 1,6-Hexanediol was added to this medium as indicated and stimulation was performed by replacing 136 mM NaCl, 2.5 mM KCl with 78.5 mM NaCl, 60 mM KCl. Unless specified otherwise, time-lapse images were acquired every 5 sec. For FRAP experiment of vesicle condensates, a single droplet was bleached by scanning with a 488 nm laser for 1 sec and fluorescence recovery was subsequently imaged at 5 sec intervals.

### Immunofluorescence

COS7 cells or cultured hippocampal neurons were fixed with 4% PFA in 4% sucrose containing PB for 15 min at RT and washed in PBS. After fixation, cells were incubated with blocking buffer (3% BSA and 0.2% triton X-100 in PBS) for 30 min at RT. Subsequent primary and Alexa Fluor-conjugated secondary antibody incubations were made in this buffer. Samples were finally mounted in Prolong Gold (Invitrogen).

### Proximity biotinylation

COS7 cells expressing the indicated constructs were plated, transfected, and labeled with pre-warmed biotin containing complete DMEM medium (500 μM biotin) for 15 min at 37°C. Cells were then washed five times with ice-cold PBS, and lysed with lysis buffer (20 mM Tris-HCl, pH 8, 1% triton X-100, 10% glycerol, 137 mM NaCl, 2 mM EDTA, 1 mM PMSF, 10 mM leupeptin, 1.5 mM pepstatin, and 1 mM aprotinin). Lysates were centrifuged at 14,000 g for 20 min at 4°C and the supernatants were collected. Protein concentration was estimated with the Pierce BCA protein Assay kit (Thermo Scientific, MA). Lysates containing 1.2 mg protein for sample were then incubated overnight under rotation at 4°C with streptavidin-coated magnetic beads (Pierce) pre-washed twice with lysis buffer. The beads were subsequently washed three times with lysis buffer, additional three times with PBS to remove the detergent and resuspended in 1.2ml PBS in Eppendorf tubes. 1 ml aliquots of this suspensions were then transferred to new Eppendorf tubes, the supernatants were removed, and the beads were frozen with dry ice for LC-MS/MS analysis. The remaining portions of the samples (0.2 ml) were used to verify the enrichment of biotinylated proteins. Biotinylated proteins were eluted from the beads by boiling for 10 min in 3x sample buffer (165 mM Tris pH 6.8, 17% glycerol, 5% SDS, 0.045% DTT and 0.06% bromophenol blue) supplemented with 20 mM DTT and 2 mM biotin. Eluted proteins were processed for SDS-PAGE and western blotting. To verify the occurrence of biotinylation at the expected sites, COS7 cells were incubated with 500 μM biotin for 15 min at 37°C prior to fixation, then washed five times with PBS and incubated with blocking buffer for 30 min at RT. Finally, cells were incubated with Alexa Fluor-conjugated HA and streptavidin antibodies in blocking buffer to detect localization of mini-TurboID fusion proteins and biotinylated proteins.

### Vesicle immunoisolation

Cells expressing HA tagged proteins were washed twice with PBS, scraped into ice-cold homogenization buffer (5 mM Tris, 250 mM Sucrose, 1 mM EGTA pH 7.4) and disrupted by 15 passages through a 27-gauge syringe on ice. The homogenate was centrifuged at 1,000 g for 10 min at 4°C. The supernatant was collected (PNS, postnuclear supernatants) and ultracentrifuged at 25,000 g for 20 min 4°C (or 50,000 g for 30 min 4°C) in a Beckman TLA100.3 rotor (Optima™ TLX Ultracentrifuge, Beckman). The pellet was discarded, and the supernatant was mixed with HA magnetic beads (Pierce) that had been washed twice with homogenization buffer. This mixture was incubated overnight at 4°C. Subsequently beads were washed three times with homogenization buffer, three times with PBS and frozen in dry ice for LC-MS/MS analysis.

### Mass spectrometry analyses

Affinity-purified proteins from BioID experiments and affinity purified vesicles from anti-HA immunoisolation were buffered with 200 mM HEPES (4-(2- hydroxyethyl)-1-piperazineethanesulfonic acid) pH 7.5. Disulfide bonds were reduced using 5 mM dithiothreitol (Sigma-Aldrich) at 37°C for 1 hour, followed by alkylation of cysteine residues using 15 mM iodoacetamide (Sigma-Aldrich) in the dark at room temperature for 1 hour. Excessive iodoacetamide was quenched using 10 mM dithiotheritol. Protein mixtures were diluted in 1:6 ratio (v/v) using ultrapure water prior to digestion using sequencing grade trypsin (Worthington Biochemical) at 37°C for 16 hours. Digested peptides were subsequently desalted using self-packed C18 STAGE tips (3M Empore™)^48^ for LC-MS/MS analysis. Desalted peptides were resolubilized in 0.1% (v/v) formic acid and loaded onto HPLC-MS/MS system for analysis on an Orbitrap Q-Exactive Exploris 480 (Thermo Fisher Scientific) mass spectrometer coupled to an FAIMS Pro Interface system and Easy nanoLC 1000 (Thermo Fisher Scientific) with a flow rate of 300 nl/min. The stationary phase buffer was 0.1% formic acid, and mobile phase buffer was 0.1% (v/v) formic acid in 80% (v/v) acetonitrile. Chromatography for peptide separation was performed using increasing organic proportion of acetonitrile (5-40% (v/v)) over a 120 min gradient) on a self-packed analytical column using PicoTip™ emitter (New Objective, Woburn, MA) using Reprosil Gold 120 C-18, 1.9 μm particle size resin (Dr. Maisch, Ammerbuch-Entringen, Germany). The mass spectrometry analyzer operated in data independent acquisition mode at a mass range of 300 - 2000 Da, compensation voltages of -50/-70 CVs with survey scan of 120,000 and 15,000 resolutions at MS1 and MS2 levels, respectively. Mass spectrometry data were processed by Spectronaut™ software version 15 (Biognosys AG)^49^ using directDIATM analysis using default settings, including: oxidized methionine residues, biotinylation, protein N-terminal acetylation as variable modification, cysteine carbamidomethylation as fixed modification, initial mass tolerance of MS1 and MS2 of 15 ppm. Protease specificity was set to trypsin with up to 2 missed cleavages allowed. Only peptides longer than seven amino acids were analyzed, and the minimal ratio count to quantify a protein was 2 (proteome only). The false discovery rate (FDR) was set to 1% for peptide and protein identifications. Database searches were performed against the UniProt Chlorocebus sabaeus (Green monkey) database containing 19,232 entries (December 2021). High precision iRT calibration was used for samples processed using the same nanospray conditions^50^. Protein tables were filtered to eliminate identifications from the reverse database and also common contaminants. The SynGO enrichment tool^40^ was used to analyze the proteins that enriched in synaptophysin-HA and ATG9A-HA samples.

### Statistical analysis

Data are presented as mean ± SD or SEM, with n indicating the number of independent experiments. Statistical significance was determined using the Student’s two-sample *t* test or paired *t* test for the comparison of two independent groups. ANOVA followed by Tukey’s honest significant difference (HSD) post hoc test was applied for multiple comparisons. All images were analyzed with ImageJ and Sigma plot, Origin 9.0 and Sigma plot were used for statistical comparisons.

## Acknowledgments

We thank Drs. Wade Harper, Miguel Gonzalez-Lozano, Tobias Walther and Zon Weng Lai (Harvard) for help and discussion. This research was supported in part by grants from the NIH (NS36251 and DA018343) and the Parkinson Foundation (PF-RCE-1946) to PDC and a fellowship from the National Research Foundation of Korea (2019R1A6A3A03031300) to DP.

## Author contributions

D.P. and P.D.C. designed the experiments. Y. W. and A. B. performed CLEM experiments and LC-MS/MS, respectively, in collaboration with D.P., and D.P. performed everything else. D.P. and P.D.C. wrote the paper. All authors read and approved the final manuscript.

## Competing interests

The authors declare no competing interests.

---

**Extended Data Fig. 1.**
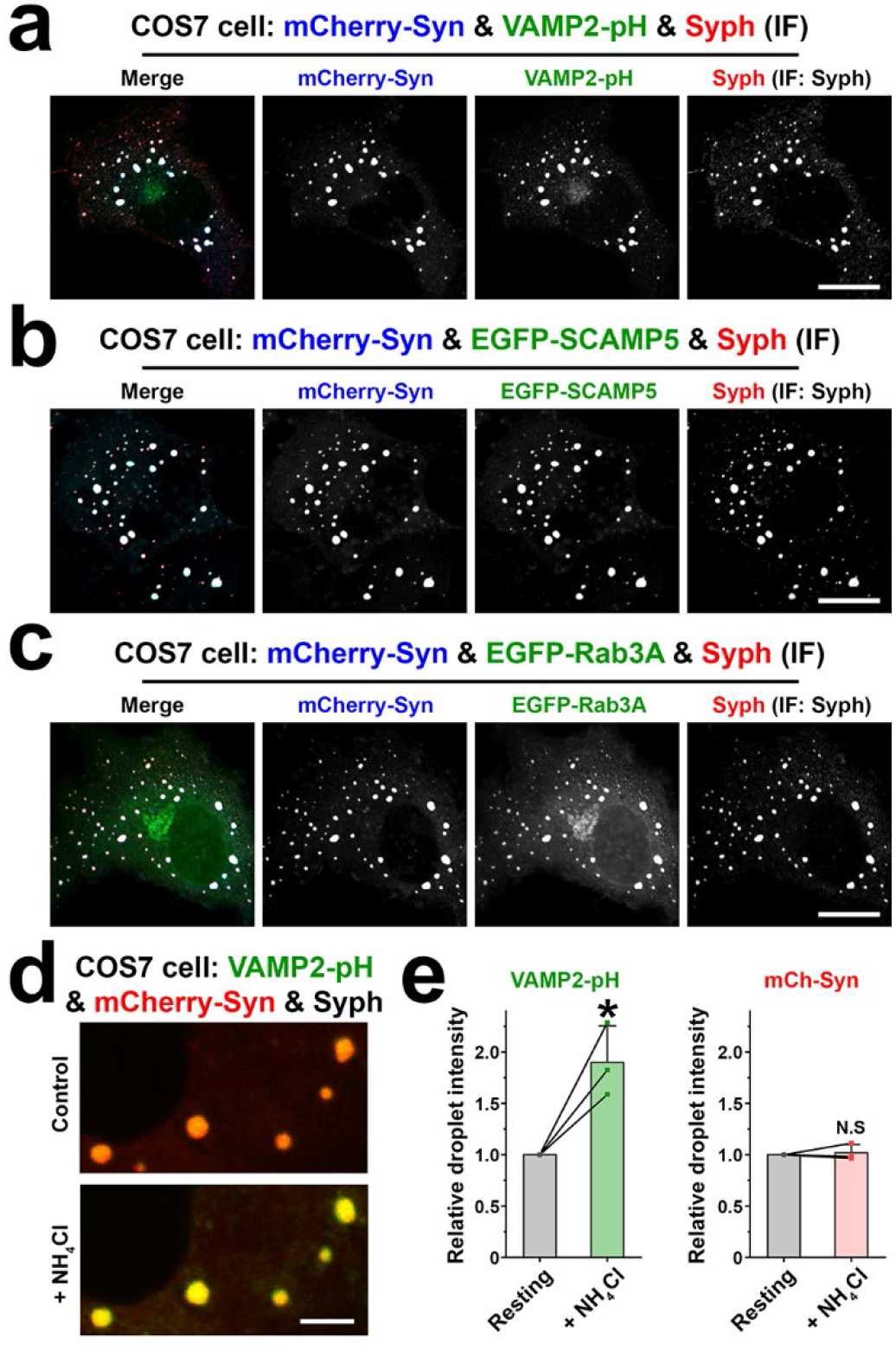
SV proteins coassemble into droplets formed by synaptophysin and synapsin. a-c, COS7 cells were transfected as indicated and synaptophysin was detected by immunofluorescence using anti-synaptophysin antibodies. d, COS7 cell expressing synaptophysin, VAMP2-pH and mCherry-synapsin before and after treatment with NH_4_Cl to alkalinize the acidic lumen of the vesicles and thus increase pHluorin fluorescence. e, Quantification of the fluorescence changes after the addition of NH_4_Cl. Values are means ± SD; N.S., not significant; \**p* < 0.05 by paired t test (n = 3 independent experiments). Scale bars, a-c = 20 μm, d = 5 μm.

**Extended Data Fig. 2.**
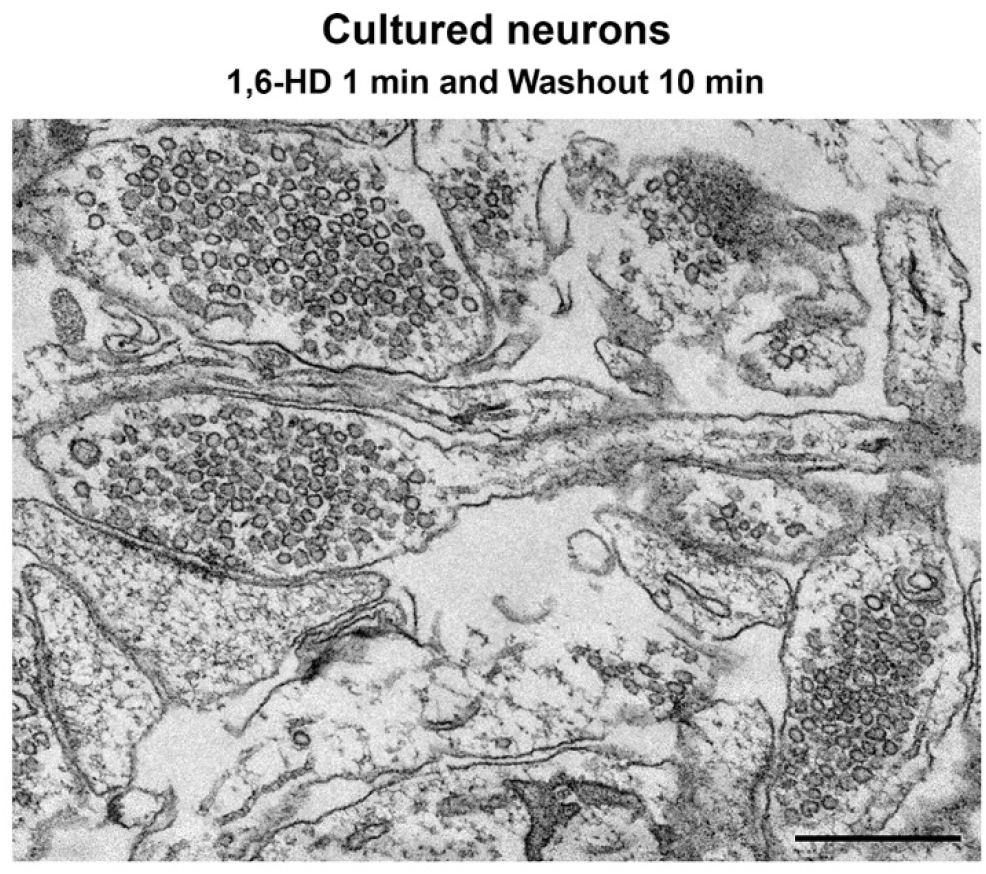
Reversible dispersion of SV clusters by 1,6-Hexanediol in nerve terminals. Cultured hippocampal neurons were washed for 10 min after 1 min incubation with 1,6-Hexanediol and then fixed for transmission electron microscopy (TEM). Scale bar = 500 nm.

**Extended Data Fig. 3.**
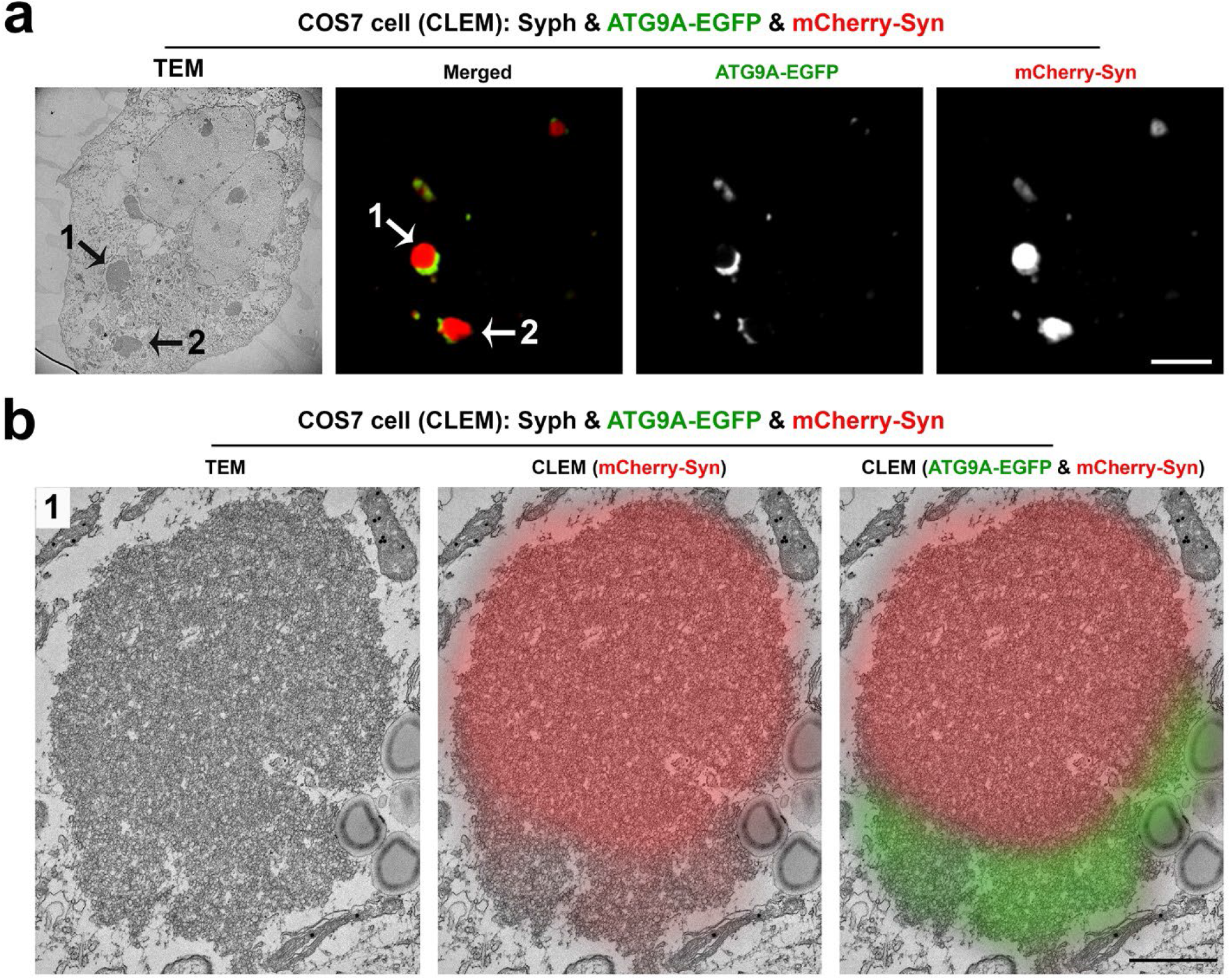
The synapsin phases include two distinct vesicle clusters positive for synaptophysin and ATG9A, respectively. a, COS7 cells co-expressing synaptophysin, ATG9A-EGFP and mCherry-synapsin were fixed and processed for correlative light-electron microscopy (CLEM). b, High magnification of fluorescence and TEM image of droplet #1. Scale bars, a = 10 μm, b = 500 nm.

**Extended Data Fig. 4.**
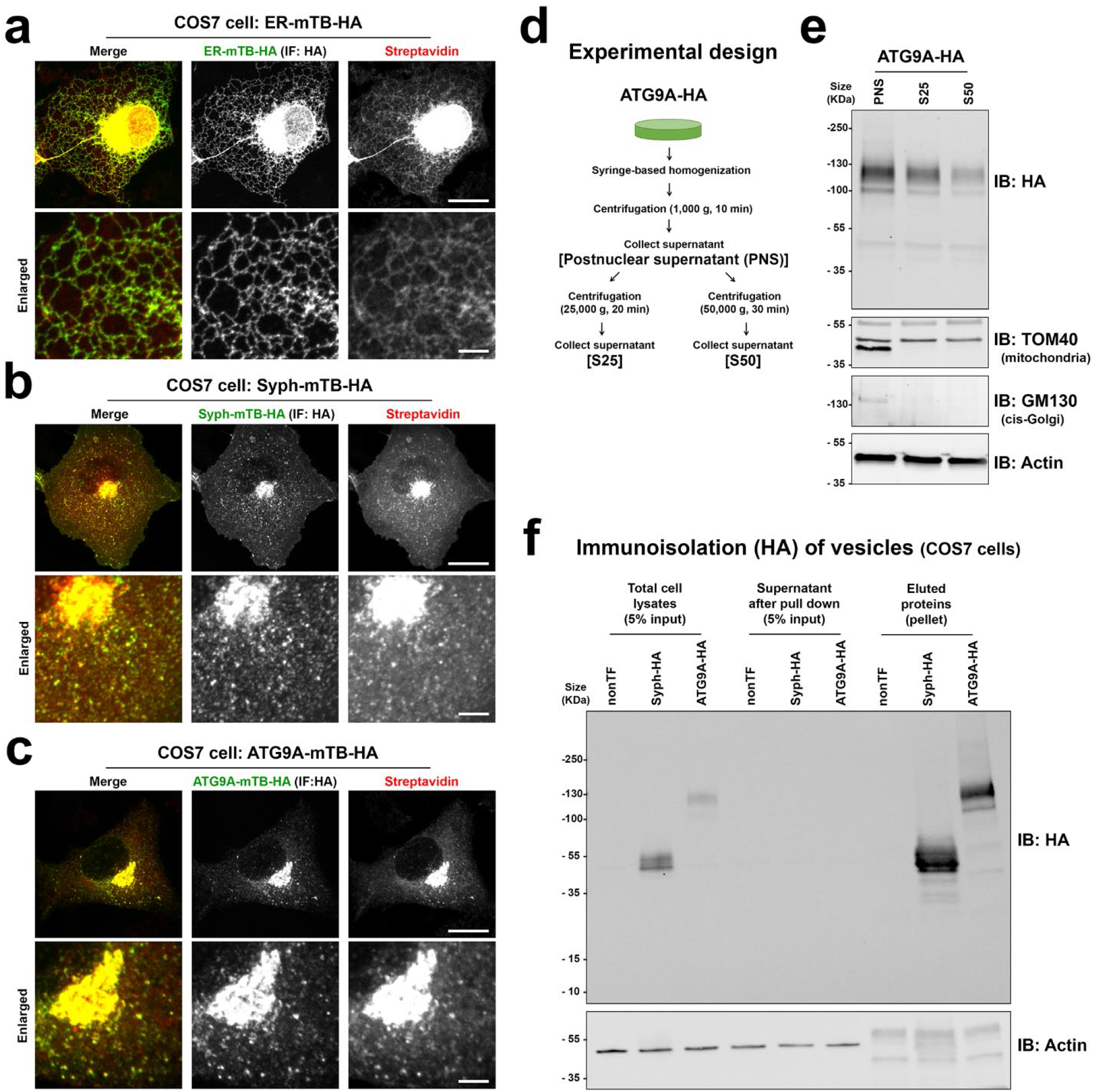
Control experiments for proteomic analysis. a-c, COS7 cells expressing ER-miniTurboID- HA (a), synaptophysin-miniTurboID-HA (b) or ATG9A-miniTurboID-HA (c) were treated with 500 μM biotin for 15 min at 37°C, then fixed and processed for anti-HA immunofluorescence or streptavidin labeling. d, Experimental design for vesicle immunoisolation. e, Preparation of vesicle-enriched fraction for vesicle immunoisolation. The postnuclear supernatant and the supernatants resulting from two different centrifugations were analyzed by Western blotting for the indicated proteins. The S25 supernatant was chosen for the immunoisolation given its loss of Golgi and mitochondrial markers and retention of the bulk of small vesicles (ATG9A) as detected by Western blotting. f, Starting S25 fraction, and material bound and not bound by anti-HA magnetic beads were analyzed by Western blotting. Scale bars, a-c = 20 μm (5 μm for the enlarged images).

**Extended Data Fig. 5.**
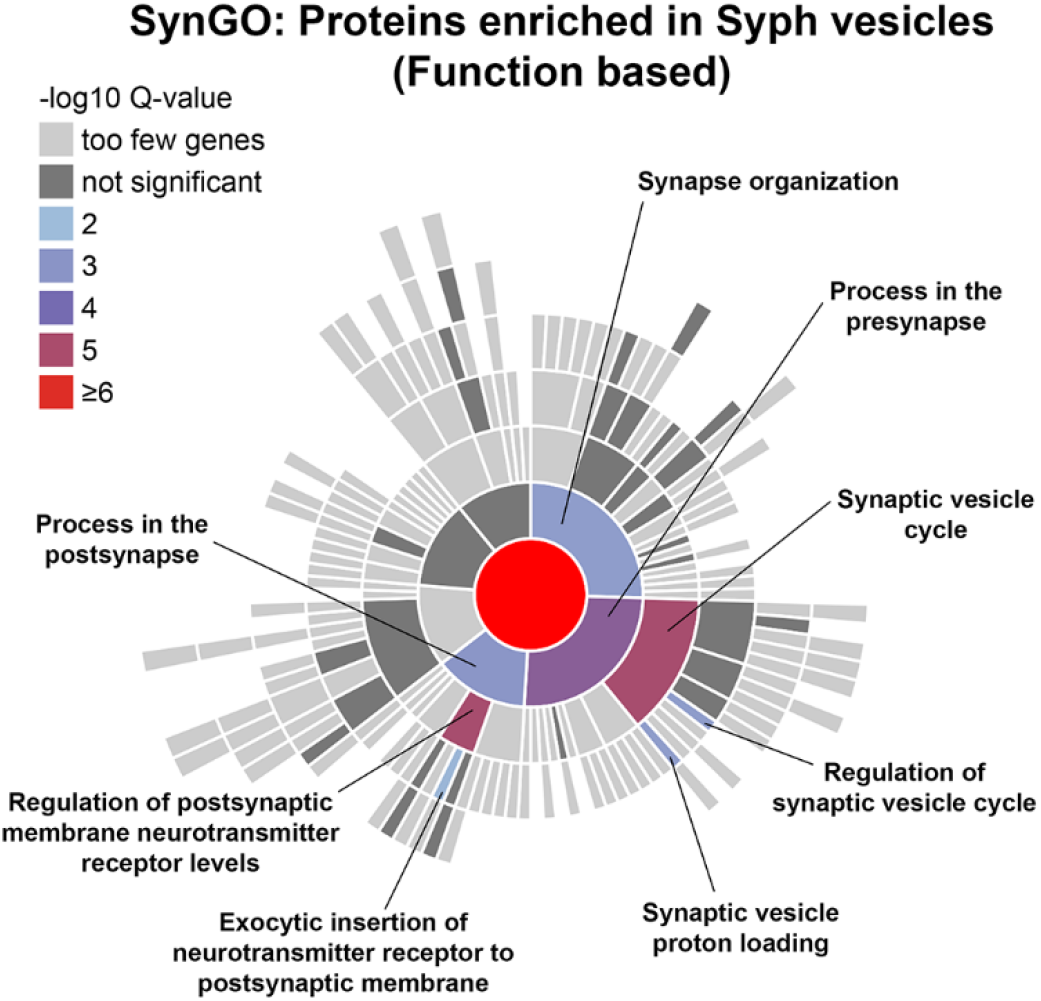
SynGO analysis of proteins enriched in synaptophysin-HA immunoisolated samples. Sunburst plot showing synaptic function annotations of the proteins enriched in synaptophysin-HA versus ATG9A- HA immunoisolated fractions. Inner rings are parent terms of more specific child terms in the outer rings. Colors represent enrichment Q value (-log10 values).

**Supplementary Video 1**. 1,6-Hexanediol (1,6-HD) dependent reversible dispersion of the condensates formed by synaptophysin, VAMP2-pH (green) and mCherry-synapsin (red) coexpression. Related to Fig. 2e.

**Supplementary Video 2**. 1,6-Hexanediol (1,6-HD) dependent reversible dispersion of the SV clusters in cultured neurons. Neurons were cotransfected with EGFP-SCAMP5 (an integral SV membrane protein, green) and mCherry- synapsin (a SV associated protein, red). Related to Fig. 2f-i.

**Supplementary Video 3**. SV exocytosis, as reflected by the increase in vGlut-pHluorin fluorescence, in the presence or absence of 1,6-Hexanediol (1,6-HD). vGlut-pHluorin and mCherry-synapsin expressing neurons were treated as indicated. Related to Fig. 2k,l.

**Supplementary Video 4**. 1,6-Hexanediol (1,6-HD) dependent reversible dispersion of two distinct vesicle clusters formed by synaptophysin, ATG9A-EGFP and mCherry-synapsin coexpression in COS7 cells. Related to Fig. 4f.

**Supplementary Video 5**. Z stack images of COS7 cells expressing ATG9A^Y8A^. The cis-Golgi was detected by immunofluorescence using GM130 antibodies. 0.2 μm / each Z sections (total 42 images). Related to Fig. 7b.

## Notes

### Competing Interest Statement

The authors have declared no competing interest.

